# Color is necessary for specialized face learning in the Northern paper wasp, *Polistes fuscatus*

**DOI:** 10.1101/2021.10.03.462925

**Authors:** Christopher M. Jernigan, Jay A. Stafstrom, Natalie C. Zaba, Caleb C. Vogt, Michael J. Sheehan

## Abstract

Visual individual recognition requires animals to distinguish among conspecifics based on appearance. Though visual individual recognition has been reported in a range of taxa, the features that animals rely on to discriminate between individuals are often not well understood. Northern paper wasp females, *Polistes fuscatus*, possess individually distinctive color patterns on their faces, which mediate individual recognition. It is currently unclear what facial features *P. fuscatus* use to distinguish individuals. The anterior optic tubercle, a chromatic processing brain region, is especially sensitive to social experience during development, suggesting that color may be important for recognition in this species. We sought to test the roles of color in wasp facial recognition. Color may be important simply because it creates a pattern. If this is the case, then wasps should perform similarly when discriminating color or grayscale images of the same faces. Alternatively, color itself may be important for recognition, which would predict poorer performance on grayscale image discrimination relative to color images. We found wasps trained on grayscale faces, unlike those trained on color images, did not perform better than chance. Suggesting that color is necessary for the recognition of an image as a face by the wasp visual system.

## INTRODUCTION

Individual recognition requires that animals discriminate among individuals within a population that share many features [1, 2]. Reliably recognizing individuals of the same age-class and species, who tend to share many similar traits, poses a challenge for animals. One solution to this problem is for animals to evolve more individually distinctive traits that facilitate individual identification. Evidence for identity signal elaboration has been found in visual [3–5], acoustic [6, 7], and chemical modalities [8, 9] across a range of taxa. Additionally, perceptual adaptations among receivers can provide improved discrimination of identity cues and signals [10–13]. Despite evidence for perceptual adaptations for individual recognition, we have little understanding of which aspects of identity cues or signals animals use to discriminate among individuals. There are often many features that vary and could be used as aspects of identity signals, but it is unclear which are relevant to discrimination. Understanding which features are used for individual recognition will result in two advances. First, uncovering which aspects of variable features contribute to identity will more clearly define the perceptual and cognitive processes by which animals recognize individuals. Second, understanding which features are most important for recognition will provide insights into the evolutionary pressures shaping identity signal diversity.

The greatest progress toward understanding the features encoding identity has been made in primates, for which facial recognition is achieved by identifying small differences in facial morphology and the relative position of internal facial features [4, 14, 15]. In primates, the detection of these facial identities is pattern dependent and does not rely on color—i.e. we can recognize faces in greyscale [15–17]. However, color may play a role in face detection and segmentation from the background environment [18–20]. The visually mediated individual recognition system within primates has provided interesting insights into how visual identities are processed and the constraints on what is required for the nervous system to recognize a stimulus as a face.

Facial individual recognition analogous to primates has independently evolved in paper wasps. Females of the Northern paper wasp, *Polistes fuscatus*, females possess highly diverse and individually distinctive colored face patterning [21]. These facial patterns alone can be used by female wasps to discriminate among nest mates or images [12, 22]. In fact, *P. fuscatus* female wasps are specialized for learning face stimuli, and their ability to discriminate among images is dramatically reduced when images are digitally altered, either by removal of the antenna or re-arrangement of the internal features of the face [12]. This finding and others suggests that like primates, faces in *P. fuscatus* are holistically processed [23] and seem to be special perceptual objects [12].

*P. fuscatus* facial patterns typically consist of only 3 pigments: a reddish-brown pigmentation, a yellow pigmentation, and a melanized black pigmentation [24, 25]. Most if not all wasp faces could be discriminated against using achromatic patterning information alone, simply using edge detection and luminance in the long-wavelength visual channel to identify the spatial relations of the various pattern features. Thus, the colorful identity signals in wasps could have evolved for the purpose of providing contrast to make patterns that are processes achromatically. Tibbetts, Desjardins [26] found that specialized facial discrimination abilities require early experience with conspecifics. Using a similar isolation assay we recently found that individuals reared in isolation also have significantly less relative brain investment in a chromatic processing center, the anterior optic tubercle [27]. This finding suggests that color in addition to spatial patterning may be critical for face discrimination in *P. fuscatus*.

Here we tested animals in an operant conditioning assay using either color images of wasps or grayscale versions of those same images to test the role of color in facial recognition in *P. fuscatus*. On one hand, color may be important simply because it creates a pattern. If this is the case, then wasps should perform similarly when discriminating color or grayscale images of the same faces. Alternatively, color itself may be important for recognition that an image is a face, which would predict poorer performance on grayscale discrimination relative to color images.

## METHODS

*Polistes fuscatus* gynes were collected in Ithaca and Erin, NY in Aug. and Sept. 2020. After collection, wasps were housed individually in small cups in the laboratory and provided with a natural light cycle. Each wasp was provided with construction paper and *ad libitum* sugar and water.

### Training stimuli and apparatus

We first selected images of 6 paper wasps with distinct facial patterns and standardized contrasts in Adobe Photoshop (CC2020). We then selected the internal facial patterns from each image and placed them onto the same wasp image with a standard gray background, creating 6 unique wasp images with the only features that vary being the facial patterns (body, antennae, mandibles, and eyes are identical across face images). Grayscale versions of all standardized images were then created using the grayscale function in Photoshop (Figure 1a), generating a total of 12 standardized images (6 color and 6 grayscale). Images were sized to 1.3 cm x 0.6 cm and printed on Canon Photo Paper Plus Glossy II using a Canon Pixma iP110 inkjet photo printer (1200 DPI). Full resolution training image files are available upon request.

**Figure 1.**
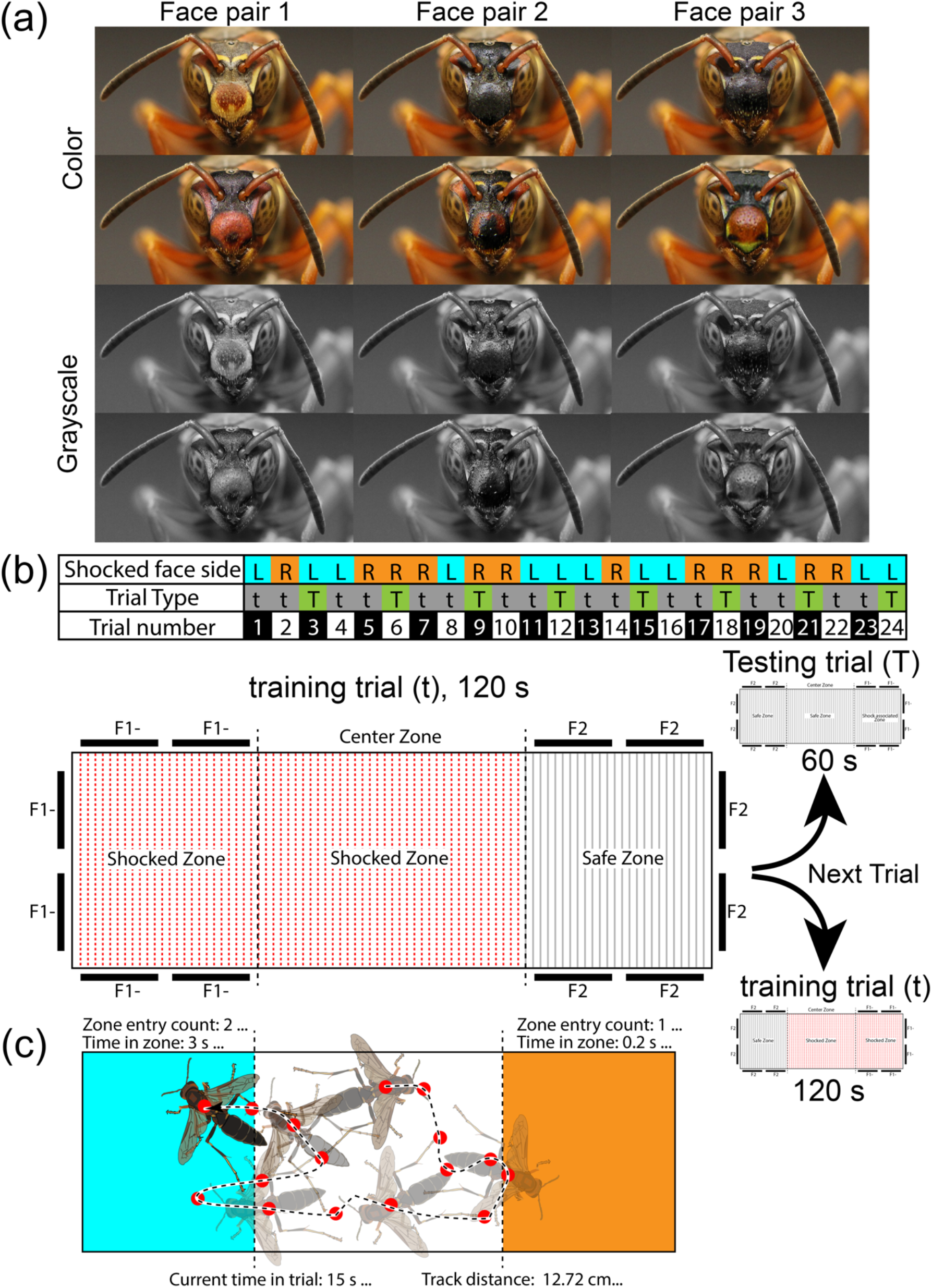
Stimuli and training paradigm used for this study. (a) Image set used in this study. A total of 6 unique wasp stimuli with identical antenna, mandible positions, and body backgrounds were used to test face discrimination in Polistes fuscatus. Each wasp stimuli also had a corresponding grayscale image used to train half of the animals. (b) Schematic of pseudorandomized side pairing of shock associated stimulus over trials, including side switching of shock associated stimulus and trial type. Animals experienced two trial types: training (t) and Testing (T). On training trials wasps were allowed to freely move about the arena for 120 seconds and were shocked in the central every 10 seconds and when entering the shock associated stimulus zone every 3 seconds. Every third trial was a Testing trial, in which wasps were allowed to freely move about the arena for 60 seconds and no shocks were delivered. To scale schematic of rectangular training arena schematic with electrical grid flooring, acrylic sheet cover, and removable central zone barriers. Red highlighted sections are electrified. Two stimuli images were affixed to each wall of the choice zones denoted by dark rectangles and faces were denoted by F1 and F2 respectively, (−) denotes electrical shock pairing. (d) Diagram of animal tracking methods and data acquisition. Red dot denotes tracked position on the wasp. Distance of track, left and right side zone entries, and time spent in each zone were collected during each trial.

We 3D printed identical rectangular arenas with inner dimensions of, 3.6 cm x 10.7 cm x 7 mm. Each arena had two sets of stimulus images affixed to all 3 walls, making up each end zone of the arena: 6 images of one face at one end and 6 images of a second face at the opposite end (Figure 1b). Arenas were placed on a printed copper electrical grid (Figure 1b). A 4.0V mild electric shock was manually delivered using a variac transformer through the grid on the floor of the arena. This level of shock was mild enough to not harm or kill the wasp, but strong enough to visibly see the wasp react when a shock was delivered [12]. A clear acrylic sheet was placed over the arena to contain the wasp and ensure the wasp always contacted the electrical grid during training. The arena was illuminated equally using three overhead LED lights. Trials were recorded using an iPhone (Apple, iPhoneX) or Go-pro camera (Hero4 Black).

### Training paradigm

Each wasp was trained to discriminate between one pair of face images, either in color or grayscale, and received 16 training trials and 8 testing trials (Figure 1b). Each training trial lasted 120 seconds and each testing trial lasted 60 seconds (Figure 1b). During training trials, a mild electric shock was associated with one of the two stimulus images. On testing trials no electric shock was delivered to the animal. The side of the arena that had the shocked-paired face was switched in a pre-determined pseudorandomized pattern across trials and each wasp experienced the shock-associated face an equal number of times on the left and right sides of the arena (Figure 1b). Prior to each trial, wasps were confined in the middle of the arena by two clear removable barriers. Once the wasp was acclimated to the arena a trial would begin and the barriers were removed, at which point wasps were allowed to freely move around the arena.

During training trials if the wasp did not move for 10s after being placed in the center area of the arena, a single shock was delivered. Each time the wasp thorax (or >50% of the body) crossed the boundary into the zone of the arena containing the shock-associated face, a mild shock was manually delivered every 3 seconds until the wasp exited the zone. After 120 seconds in the arena, the wasp was removed from the arena and placed in a holding container with sugar and water for 1 minute. On testing trials, the wasp was placed in the center zone of the arena until acclimated and barriers were then removed. The wasp was then allowed to explore the arena for 60 seconds without being shocked. Again after 60 seconds the wasp was removed from the arena and placed in a holding container with sugar and water for 1 minute.

Each set of trials consisted of two training trials followed by a single testing trial (Figure 1b). Between each trial, the apparatus was wiped with water to remove or homogenize possible chemical cues released by wasps. Between each individual wasp, the training apparatus was wiped with ethanol. Training was conducted in a windowless room, where the only source of light was the LED lights evenly illuminating the arena.

### Data metrics and analyses

Deep Lab Cut [28, 29] was used to track each wasp’s thorax (as the thorax was used as the point of reference to induce shock) throughout the trial. we labeled 589 frames taken from 89 videos (then 95% of the labelled frames were used for training). We used a ResNet-50-based neural network with default parameters for 500,000 training iterations. We found the test error was: 2.21 pixels, train: 1.64 pixels. We then used a p-cutoff of 0.9 to condition the X,Y coordinates for future analysis. We then used SimBA [30] to extract the movement distance, average velocity, and proportion of time spent in each of the stimulus zones of the arena (shocked and safe zones, Figure 1c). Additionally, first entry side and the latency to first entry in the non-shock-associated face side (safe zone) was manually scored from each video. To assess learning performance between wasps trained to color or grayscale faces we used a chi-squared test using the total number of correct and incorrect choices over the testing trials compared to random chance [31, 32]. We also used mixed effect models to test differences between treatments using continuous data with treatment (color vs. grayscale stimuli), the trial number, the previously shocked side, and their interactions as fixed effects and wasp ID and trainer as a random effects using the lme4 function in R [33, 34].

## RESULTS

We trained a total of 17 wasps using color images and 16 wasps using grayscale images. All other training protocols were identical (Figure 1b). On testing trials, we found that wasps trained using color faces chose to first enter the safe associated face zone (hereafter referred to as ‘safe’ zone) significantly more than predicted by chance (X^2^= 7.1, df=1, p=0.008, Figure 2a). However, wasps that were trained using grayscale versions of the same face stimuli did not show a stimulus first-choice preference (X^2^= 0.019, df=1, p=0.891). Additionally, on testing trials the latency to enter the safe zone was significantly faster in wasps trained to color faces and treatment (color vs grayscale) was the only significant factor for this measure (t-value= 2.476, p=0.013, Figure 2b). Interestingly, on testing trials we also observed significantly more trials in which animals did not make a stimulus zone choice and instead stayed in the center zone for the full 60 unreinforced seconds when wasps were trained using grayscale faces than those trained using color faces (gray: 21 of 128 trials, color: 8 of 136 trials, X^2^= 6.431, df=1, p=0.011).

**Figure 2.**
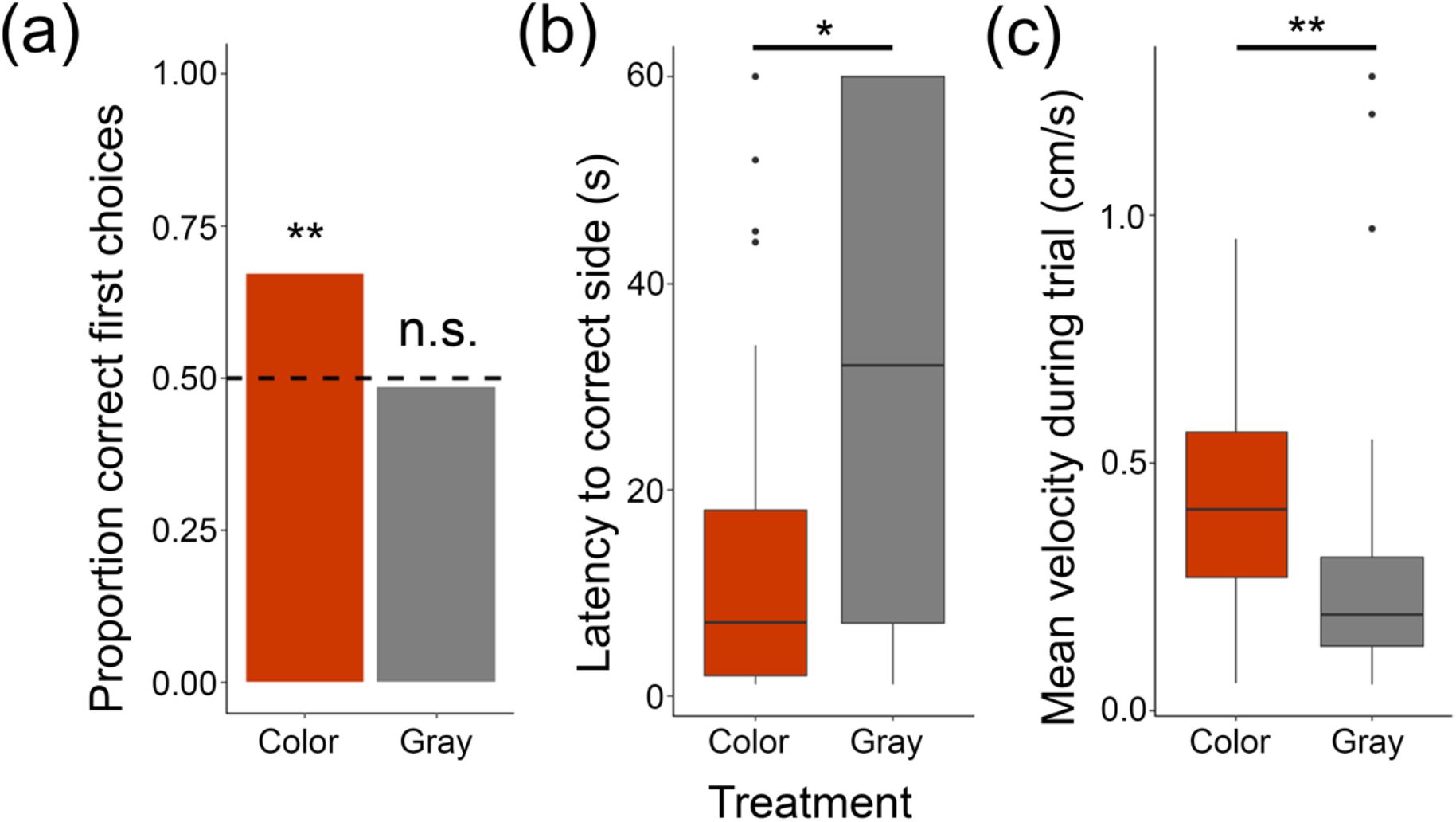
Testing trial performance of wasps trained across stimulus treatments: color (orange) or grayscale (gray). (a) Total proportion of safe associated stimulus zone first choices for all testing trials for animals trained across treatments. (b) Latency to enter safe associated stimulus zone for animals trained across treatments. (c) Averaged mean velocity (cm/s) for animals trained across treatments. Treatments denoted by coloration. Symbols denote significant differences: ** p<0.01, *p<0.05, n.s. = non-significant. See methods and results for statistical test details.

Using computer vision software, we tracked wasps and calculated distance traveled, average velocity, and proportion of time spent in each zone of the arena. On testing trials wasps trained using color faces moved longer distances at a faster mean velocity than wasps trained using grayscale faces (distance: t-value= −3.068, p=0.002; mean velocity: t-value=−3.072, p=0.002, no other factors predicted measures, Figure 2c). This could explain the latency differences between treatments, however this cannot explain the first-choice differences. Surprisingly on testing trials we also found that the proportion of time spent in the correct zone was unaffected by treatment (t-value=-1.313, p=0.189), instead the proportion of time spent in a zone was only predicted by the side shocked in the previous trial (t-value=2.286, p=0.022).

On training trials, neither treatment performed better than chance based on first-choice decisions (color: X^2^= 0.068, df=1, p=0.795; grayscale: X^2^= 0.008, df=1, p=0.929, Figure Sa) and no factor significantly affected latency to enter the non-shock associated face zone on training trials (p>0.05, Figure Sb). We also observed no difference in the number of trials in which animals did not leave the center-shocked zone between treatments on training trials (gray: 1 of 266 trials, color: 6 of 272 trials).

On training trials, we found that distance and mean velocity were unaffected by treatment (t-value= −1.653, p=0.098, Figure Sc) and instead trial number was the only factor that significantly affected the proportion of time spent in the correct zone during reinforced training trials (t-value=0.271, p=0.005, Figure Sd). Also in training trials, trial number was the only factor to significantly impacted the distance and velocity at which wasps moved (t-value=-2.950, p=0.003, Figure Se) with wasps moving slower as wasps learned the assay and trials increased. Together these data suggest that regardless of treatment wasps changed their behavior consistently over trials, but only wasps trained using color stimuli performed better than chance on the assay during unreinforced trials.

## Discussion

Facial recognition in the paper wasp is chromatic-dependent – *Polistes fuscatus* readily discriminated between different wasp face images when color is present but did not discriminate between face images rendered in grayscale (Figure 2). Wasps trained with colored faces made more correct decisions during testing trials and made these decisions more quickly. Wasps trained with color faces were also generally more active during testing trials, both in terms of body movement velocity and total distance travelled. Together, our results support the hypothesis that color is necessary for facial recognition in *Polistes fuscatus* and that images of wasps faces in grayscale are treated differently from that of colored faces. We propose that color is an important component to the “holistic faceness” required for facial recognition in paper wasps.

### Chromatic-dependent facial recognition

In humans and non-human primates, facial processing and recognition is not dependent on color [15–17]. Primates can discriminate between faces when presented greyscale faces of conspecifics, much like how we presented grayscale images in our behavioral assay [16, 17]. Primate abilities of facial recognition do not depend on color, but rather depend on detecting differences in spatial relationship of critical facial features across conspecifics [15]. Wasps utilize unique color patterns as signals of identity [3]. Prior to this study whether color itself was important in wasp facial recognition was unknown. Wasps could utilize the patterns created by the color, but not require the color per se (i.e. shapes created by the color could be the necessary part of the signal, independent of which colors are used). However, our recent neuroanatomical study predicted that color may play a key role in social processing in *P. fuscatus* [27], and indeed this appears to be true at least for face acquisition. Additional tests will be required to assess if wasps use all patterning/color features equally when making discriminations. Additionally, it remains a possibility that after a face is learned achromatic patterning alone could then be sufficient for wasps to make discriminations as nests are sometimes found in dark crevices for which chromatic information would be reduced.

This study adds to a growing body of literature examining what constitutes a ‘face’ for *P. fuscatus* wasps. Thus far we know that the correct positions of internal features are required [12] and antennae or antennae and body are needed to be present [12, 23]. This study adds that facial color patterning alone can confer identity as we controlled all other aspects of body position within the images. Most importantly, this study demonstrates that chromatic information is required facial learning in *P. fuscatus*. As in primates, faces in the *P. fuscatus* system appear to be special, holistically processed perceptual objects [23, 35, 36]. While the neural mechanisms of face recognition remain elusive in *Polistes fuscatus*, the anterior optic tubercle and its downstream targets remain prime candidates due to the fact that it is socially plastic [27] and knowledge from other insects of its role in chromatic information processing [37], figure-ground detection [38] and female-female aggression [39].

#### Training paradigm and animal tracking

A major strength of the presented behavioral assay is the confirmation that unreinforced trials show robust learning performance. Unlike in prior studies of learning in *P. fuscatus* [12, 23, 32] in this study we added unreinforced testing trials every 3 trials during training, in which animals received no shock. These un-reinforced trials allow animals to display choices that are potentially unbiased by shock-escape responses. Notably we did not observe significantly increased performance over trials in first choice data in either the training or testing trials. In the case of the testing trials, though this may be an issue of statistical power combined with already high performance on the first testing trial (first test trial: 59% correct choice in color group, 7^th^ test trial: 76% correct in color group). The lack of improvement during training trials is more puzzling. Prior studies with only reinforced trials show clear increased first choice performance over successive trials across a range of stimuli [12]. Given the pattern of two training and then one testing trial, it is possible that wasps show rapid extinction after testing trials that is recovered during training leading to observed patterns. Our selected training protocol cannot at present disentangle these possibilities. Regardless, on unreinforced trials animals show clear patterns of stimulus preference. By including unreinforced trials, future work in this system will be able to assess perceptual similarity among presented stimuli [40] or identify the salient components of multimodal stimuli [41] as is common in Probiscis Extension Responses assays in other insects. The inclusion of unreinforced trials will also allow testing of additional phenomenon commonly measured in conditioning assays such as short- and long-term memory [42] , extinction [43], and the ability to conduct neural manipulations that assess memory locations [44, 45].

By using computer vision software, we tracked animal body segment positions during testing and training from video recordings. This ability opens a host of new metrics we can use to assess performance in operant conditioning assays in this system. One interesting finding is that animals trained on color stimuli moved significantly more than animals trained on grayscale stimuli. Movement alone cannot explain the increased performance of color-trained wasps. While tracking animals we only measured time spent in each area of the arena and we did not measure the precise sequence of moments at stimuli or at borders of shocked and non-shocked zones in the arena, which could shed light on other behavioral or postural differences exhibited by animals as they approached stimuli. Additional assays and more fine grain analyses of movement sequences will be required to identify potential causes of the movement pattern differences between groups. However, if color images are recognized as “wasps” and grayscale images are not, response to the perceived presence of other individuals might explain this movement difference. Generally, wasps move more in behavioral assays when interacting with a social partner than when in isolation (personal observation). It is also possible that this could also be a form of ‘learned helplessness’ in the animals which cannot discriminate images in the assay.

## Conclusion

Color is necessary for face discrimination in *Polistes* fuscatus. This work builds upon other recent findings that are beginning to define what is required for individual recognition in this species and suggests that color processing centers of the brain are likely to be important in specialized face processing.

## Declarations

### Funding information

This work was supported by NIH Grant DP2-GM128202 awarded to MJS.

### Competing interests statement

Authors declare no conflicts of interest.

### Author contributions statement

CMJ, JAS, and MJS conceived of and designed the study. JAS and NCZ trained animals. CCV aided JAS and NCZ in collecting tracking information. JAS, NCZ, and CMJ collected data from videos. CMJ and MJS analyzed data. CMJ, JAS, NCZ, and MJS wrote the manuscript.

### Ethics statement

All work was conducted in accordance with ethics requirements for invertebrate work at Cornell University.

### Consent for publication

All authors have read and approved this manuscript for publication.

---

**Supplemental Figure.**
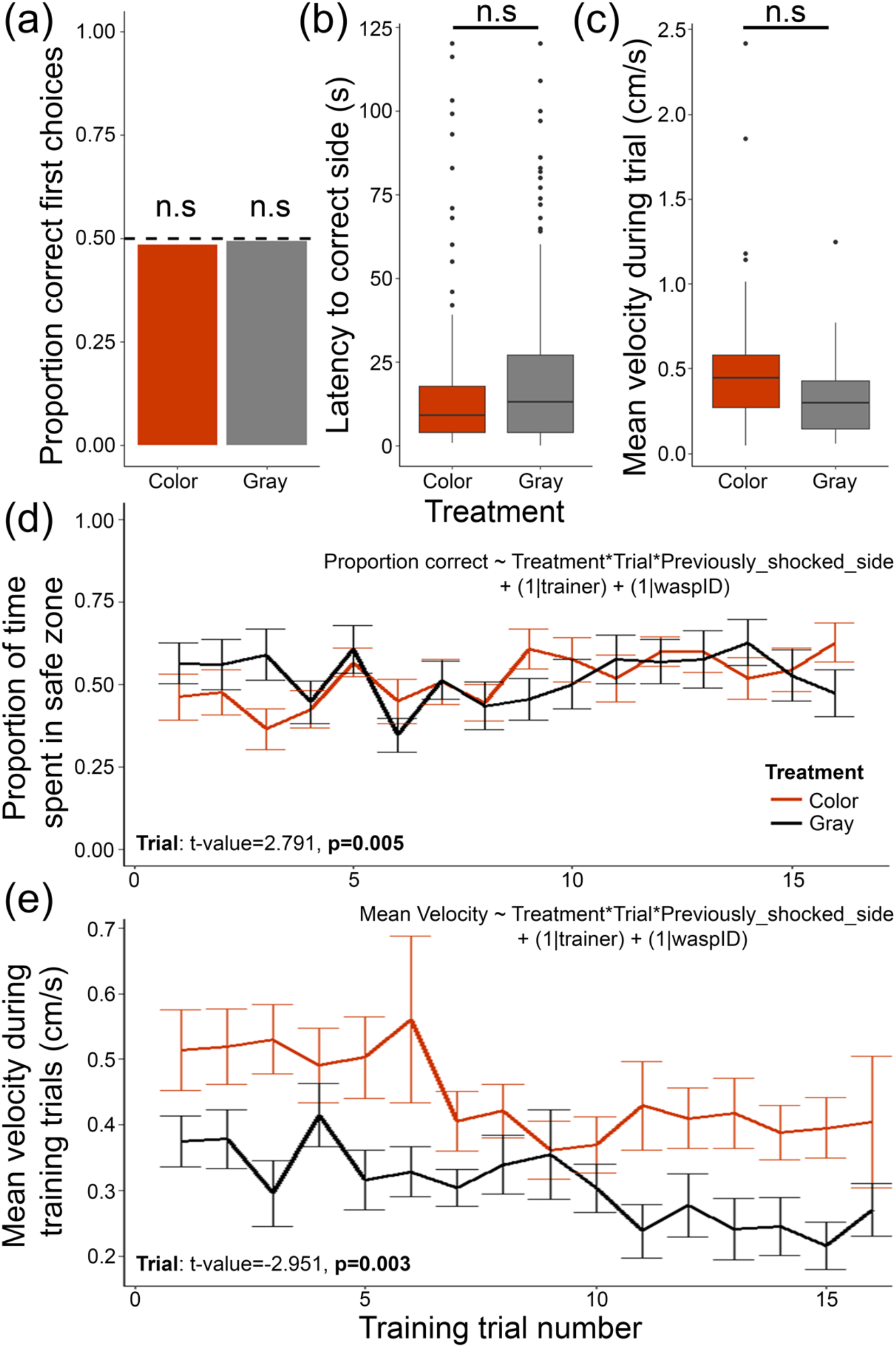
Training trial performance of wasp strained using color or grayscale stimuli. (a) Total proportion of safe associated stimulus zone first choices for all training trials for animals trained across treatments. (b) Latency to enter safe associated stimulus zone for animals trained across treatments. (c) Averaged mean velocity (cm/s) for animals trained across training treatments. (d) Proportion of time spent in the safe associated stimulus zone across training trials for animals trained in each stimulus treatment. (e) Mean velocity for animals across training trials for animals trained in each stimulus treatment. Error bars denote standard error from the mean. Symbols denote significant differences in panels (a) through (c), n.s. = non-significant. Statistical model used to compare measures in (d) and (e) are presented in each panel. Statistically significant factors are listed in the bottom of each panel; all other factors and factor interactions are non-significant. See methods and results for further statistical test details.

## Notes

### Competing Interest Statement

The authors have declared no competing interest.

## References

[1] Tibbetts, E.A. & Dale, J. 2007 Individual recognition: it is good to be different. Trends in Ecology & Evolution 22, 529–537. (doi:https://doi.org/10.1016/j.tree.2007.09.001).

[2] Tumulty, J.P. & Sheehan, M.J. 2020 What Drives Diversity in Social Recognition Mechanisms? Frontiers in Ecology and Evolution 7. (doi:10.3389/fevo.2019.00517).

[3] Sheehan, M.J. & Tibbetts, E.A. 2009 Evolution of identity signals: Frequency-dependent benefits of distinctive phenotypes used for individual recognition. Evolution 63, 3106–3113. (doi:10.1111/j.1558-5646.2009.00833.x).

[4] Sheehan, M.J. & Nachman, M.W. 2014 Morphological and population genomic evidence that human faces have evolved to signal individual identity. Nature Communications 5, 4800. (doi:10.1038/ncomms5800).

[5] Caves, E.M., Stevens, M., Iversen, E.S. & Spottiswoode, C.N. 2015 Hosts of avian brood parasites have evolved egg signatures with elevated information content. Proceedings of the Royal Society B: Biological Sciences 282, 20150598. (doi:doi:10.1098/rspb.2015.0598).

[6] Beecher, M.D. 1989 Signalling systems for individual recognition: an information theory approach. Animal Behaviour 38, 248–261. (doi:https://doi.org/10.1016/S0003-3472(89)80087-9).

[7] Pollard, K.A. & Blumstein, D.T. 2011 Social Group Size Predicts the Evolution of Individuality. Current Biology 21, 413–417. (doi:https://doi.org/10.1016/j.cub.2011.01.051).

[8] Steiger, S., Franz, R., Eggert, A.-K. & Müller, J.K. 2008 The Coolidge effect, individual recognition and selection for distinctive cuticular signatures in a burying beetle. Proceedings of the Royal Society B: Biological Sciences 275, 1831–1838. (doi:doi:10.1098/rspb.2008.0375).

[9] Sheehan, M.J., Lee, V., Corbett-Detig, R., Bi, K., Beynon, R.J., Hurst, J.L. & Nachman, M.W. 2016 Selection on Coding and Regulatory Variation Maintains Individuality in Major Urinary Protein Scent Marks in Wild Mice. PLOS Genetics 12, e1005891. (doi:10.1371/journal.pgen.1005891).

[10] Loesche, P., Higgins, B.J., Stoddard, P.K. & Beecher, M.D. 1991 Signature Versus Perceptual Adaptations for Individual Vocal Recognition in Swallows. Behaviour 118, 15–25. (doi:https://doi.org/10.1163/156853991X00175).

[11] Avilés, J.M., Vikan, J.R., Fossøy, F., Antonov, A., Moksnes, A., Røskaft, E. & Stokke, B.G. 2010 Avian colour perception predicts behavioural responses to experimental brood parasitism in chaffinches. Journal of Evolutionary Biology 23, 293–301. (doi:https://doi.org/10.1111/j.1420-9101.2009.01898.x).

[12] Sheehan, M.J. & Tibbetts, E.A. 2011 Specialized Face Learning Is Associated with Individual Recognition in Paper Wasps. Science 334, 1272–1275. (doi:10.1126/science.1211334).

[13] Hiramatsu, C., Melin, A.D., Allen, W.L., Dubuc, C. & Higham, J.P. 2017 Experimental evidence that primate trichromacy is well suited for detecting primate social colour signals. Proceedings of the Royal Society B: Biological Sciences 284, 20162458. (doi:doi:10.1098/rspb.2016.2458).

[14] Ellis, H.D., Shepherd, J.W. & Davies, G.M. 1979 Identification of Familiar and Unfamiliar Faces from Internal and External Features: Some Implications for Theories of Face Recognition. Perception 8, 431–439. (doi:10.1068/p080431).

[15] Chang, L. & Tsao, D.Y. 2017 The Code for Facial Identity in the Primate Brain. Cell 169, 1013–1028.e1014. (doi:10.1016/j.cell.2017.05.011).

[16] Rosenfeld, S.A. & Van Hoesen, G.W. 1979 Face recognition in the rhesus monkey. Neuropsychologia 17, 503–509. (doi:https://doi.org/10.1016/0028-3932(79)90057-5).

[17] Moscovitch, M., Winocur, G. & Behrmann, M. 1997 What Is Special about Face Recognition? Nineteen Experiments on a Person with Visual Object Agnosia and Dyslexia but Normal Face Recognition. Journal of Cognitive Neuroscience 9, 555–604. (doi:10.1162/jocn.1997.9.5.555).

[18] Yip, A.W. & Sinha, P. 2002 Contribution of Color to Face Recognition. Perception 31, 995–1003. (doi:10.1068/p3376).

[19] Bindemann, M. & Burton, A.M. 2009 The Role of Color in Human Face Detection. Cognitive Science 33, 1144–1156. (doi:https://doi.org/10.1111/j.1551-6709.2009.01035.x).

[20] Or, C.C.-F., Retter, T.L. & Rossion, B. 2019 The contribution of color information to rapid face categorization in natural scenes. Journal of Vision 19, 20–20. (doi:10.1167/19.5.20).

[21] Sheehan, M.J., Choo, J. & Tibbetts, E.A. 2017 Heritable variation in colour patterns mediating individual recognition. Royal Society Open Science 4, 161008. (doi:doi:10.1098/rsos.161008).

[22] Tibbetts, E.A. 2002 Visual signals of individual identity in the wasp Polistes fuscatus. Proceedings of the Royal Society of London. Series B: Biological Sciences 269, 1423–1428. (doi:10.1098/rspb.2002.2031).

[23] Tibbetts, E.A., Pardo-Sanchez, J., Ramirez-Matias, J. & Avarguès-Weber, A. 2021 Individual recognition is associated with holistic face processing in *Polistes* paper wasps in a species-specific way. Proceedings of the Royal Society B: Biological Sciences 288, 20203010. (doi:doi:10.1098/rspb.2020.3010).

[24] Enteman, W.M. 1904 Coloration in polistes, Carnegie Institution.

[25] Tumulty, J.P., Miller, S.E., Van Belleghem, S.M., Weller, H.I., Jernigan, C.M., Vincent, S., Staudenraus, R.J., Legan, A.W., Polnaszek, T.J., Uy, F.M.K., et al. 2021 Evidence for a selective link between cooperation and individual recognition. bioRxiv, 2021.2009.2007.459327. (doi:10.1101/2021.09.07.459327).

[26] Tibbetts, E.A., Desjardins, E., Kou, N. & Wellman, L. 2019 Social isolation prevents the development of individual face recognition in paper wasps. Animal Behaviour 152, 71–77. (doi:https://doi.org/10.1016/j.anbehav.2019.04.009).

[27] Jernigan, C.M., Zaba, N.C. & Sheehan, M.J. 2021 Age and social experience induced plasticity across brain regions of the paper wasp Polistes fuscatus. Biology Letters 17, 20210073. (doi:10.1098/rsbl.2021.0073).

[28] Mathis, A., Mamidanna, P., Cury, K.M., Abe, T., Murthy, V.N., Mathis, M.W. & Bethge, M. 2018 DeepLabCut: markerless pose estimation of user-defined body parts with deep learning. Nature Neuroscience 21, 1281–1289. (doi:10.1038/s41593-018-0209-y).

[29] Nath, T., Mathis, A., Chen, A.C., Patel, A., Bethge, M. & Mathis, M.W. 2019 Using DeepLabCut for 3D markerless pose estimation across species and behaviors. Nature Protocols 14, 2152–2176. (doi:10.1038/s41596-019-0176-0).

[30] Nilsson, S.R., Goodwin, N.L., Choong, J.J., Hwang, S., Wright, H.R., Norville, Z.C., Tong, X., Lin, D., Bentzley, B.S., Eshel, N., et al. 2020 Simple Behavioral Analysis (SimBA) – an open source toolkit for computer classification of complex social behaviors in experimental animals. bioRxiv, 2020.2004.2019.049452. (doi:10.1101/2020.04.19.049452).

[31] Tibbetts, E.A. & Sheehan, M.J. 2011 Facial Patterns are a Conventional Signal of Agonistic Ability in Polistes exclamans Paper Wasps. Ethology 117, 1138–1146. (doi:10.1111/j.1439-0310.2011.01967.x).

[32] DesJardins, N. & Tibbetts, E.A. 2018 Sex differences in face but not colour learning in Polistes fuscatus paper wasps. Animal Behaviour 140, 1–6. (doi:https://doi.org/10.1016/j.anbehav.2018.03.012).

[33] R Core Team. 2020 R: A Language and Environment for Statistical Computing. (R Foundation for Statistical Computing.

[34] Bates, D., Mächler, M., Bolker, B. & Walker, S. 2015 Fitting linear mixed-effects models using lme4. Journal of Statistical Software 67, 1–48. (doi:10.18637/jss.v067.i01).

[35] Tanaka, J.W. & Farah, M.J. 1993 Parts and wholes in face recognition. The Quarterly Journal of Experimental Psychology Section A 46, 225–245. (doi:10.1080/14640749308401045).

[36] Tsao, D.Y. & Livingstone, M.S. 2008 Mechanisms of Face Perception. Annual Review of Neuroscience 31, 411–437. (doi:10.1146/annurev.neuro.30.051606.094238).

[37] Mota, T., Gronenberg, W., Giurfa, M. & Sandoz, J.-C. 2013 Chromatic Processing in the Anterior Optic Tubercle of the Honey Bee Brain. The Journal of Neuroscience 33, 4. (doi:10.1523/JNEUROSCI.1412-12.2013).

[38] Aptekar, J.W., Keleş, M.F., Lu, P.M., Zolotova, N.M. & Frye, M.A. 2015 Neurons Forming Optic Glomeruli Compute Figure–Ground Discriminations in *Drosophila*. The Journal of Neuroscience 35, 7587. (doi:10.1523/JNEUROSCI.0652-15.2015).

[39] Schretter, C.E., Aso, Y., Robie, A.A., Dreher, M., Dolan, M.-J., Chen, N., Ito, M., Yang, T., Parekh, R., Branson, K.M., et al. 2020 Cell types and neuronal circuitry underlying female aggression in Drosophila. eLife 9, e58942. (doi:10.7554/eLife.58942).

[40] Guerrieri, F., Lachnit, H., Gerber, B. & Giurfa, M. 2005 Olfactory blocking and odorant similarity in the honeybee. Learning & Memory 12, 86–95.

[41] Jernigan, C.M., Roubik, D.W., Wcislo, W.T. & Riveros, A.J. 2014 Color-dependent learning in restrained Africanized honey bees. Journal of Experimental Biology 217, 337–343.

[42] Xia, S.-Z., Feng, C.-H. & Guo, A.-K. 1998 Temporary Amnesia Induced by Cold Anesthesia and Hypoxia in Drosophila. Physiology & Behavior 65, 617–623. (doi:https://doi.org/10.1016/S0031-9384(98)00191-7).

[43] Eisenhardt, D. 2012 Extinction Learning in Honey Bees. In Honeybee Neurobiology and Behavior: A Tribute to Randolf Menzel (eds. C.G. Galizia, D. Eisenhardt & M. Giurfa), pp. 423–438. Dordrecht, Springer Netherlands.

[44] Erber, J., Masuhr, T. & Menzel, R. 1980 Localization of short-term memory in the brain of the bee, Apis mellifera. Physiological Entomology 5, 343–358.

[45] Packard, M.G. & White, N.M. 1991 Dissociation of hippocampus and caudate nucleus memory systems by posttraining intracerebral injection of dopamine agonists. Behavioral Neuroscience 105, 295–306. (doi:10.1037/0735-7044.105.2.295).

